# Evaluating satellite and modeled lake surface water temperature across the contiguous United States

**DOI:** 10.64898/2026.03.06.708569

**Authors:** Blake A. Schaeffer, Hannah Ferriby, Wilson Salls, Natalie Reynolds, Jeffrey W. Hollister, Betty Kreakie, Stephen Shivers, Brent Johnson, Olivia Cronin-Golomb, Kate Meyers, Maxwell Beal

## Abstract

We developed a model to predict surface water temperature across U.S. lakes using satellite remote sensing and in situ observations to enhance cyanobacterial harmful algal bloom (cyanoHAB) forecasting. The study focused on Sentinel-3 Ocean and Land Colour Instrument (OLCI) sensor resolved lakes. We developed random forest models using both Landsat-derived and in-situ-measured surface water temperature. Landsat models offered broad spatial and temporal coverage of all OLCI resolved lakes, but they were sensitive to cloud cover and required filtering to minimize error. In contrast, the in situ model represented fewer OLCI resolved lakes, but yielded lower mean absolute error and bias. The models predicted lake surface temperature across the entire calendar year, with best performance (RMSE_applied_=1.11; bias_applied_=0.01; MAE_applied_=0.77) from the in situ model. This approach allowed for the continuous prediction of lake surface temperatures from 1.1 to 31.6 °C for unfrozen, open–water conditions critical for improving the accuracy of cyanoHAB forecasting. A key strength of this study was the use of an extensive dataset and model validation against in situ observations, which improved predictive accuracy throughout the year across all seasons. The predictive model offers a water resource tool for management, ecosystem protection, and public health.

## Introduction

Routine monitoring of lake surface water temperature has become essential to protect human health and aquatic ecosystems. Surface water temperatures within the photic zone play a critical role in lake ecosystem processes and have been reported to increase globally with changing climate (O’Reilly et al. 2015; Rose et al. 2016). Biological growth and metabolism generally increase with higher surface temperatures, promoting autotrophic production, particularly by cyanobacteria (Kraemer et al. 2017). Cyanobacteria have a competitive advantage over green algae given their ability to control buoyancy and to obtain light and nutrients more efficiently with warmer temperatures (Carey et al. 2012; Paerl and Huisman 2008; Wagner and Adrian 2009). Cyanobacteria can proliferate and form harmful algal blooms (cyanoHABs) where the biomass and cyanotoxins may reduce lake biodiversity, alter food webs, and impact human, animal, and ecosystem health (Hilborn and Beasley 2015; Huisman et al. 2018). CyanoHABs may also have adverse economic impacts resulting from beach closures and other reduced recreational opportunities. Lakes in the United States with cyanoHAB events increased 8.3% from 2007 to 2012 based on the National Lakes Assessment (NLA) (U.S. EPA 2009; U.S. EPA 2011). Drinking water sources impacted by cyanoHABs can also lead to acute and chronic health effects as well as taste and odor problems (Carmichael and Boyer 2016; Codd et al. 2005). In addition to the biological responses to temperature, physical processes such as thermal stratification impact lake turnover and nutrient cycling (Kraemer et al. 2015; Michelutti et al. 2016; Richardson et al. 2017). Given that surface water temperature may serve as a keystone variable for aquatic ecosystems and is directly linked to formation and persistence of cyanoHABs, the need for increased spatial and temporal monitoring as well as predictive modeling has received much attention (Kreakie et al. 2021; Myer et al. 2020; O’Reilly et al. 2015; Peeters et al. 2002).

Collecting lake surface temperature data across large spatial extents and over long time periods is resource intensive. Additionally, variations in sampling methods such as depth at which temperature measures were collected and sampling frequency contribute to complexities in integrating existing datasets to infer long-term changes on the ecology of aquatic systems. There are several examples of long-term, high-frequency lake temperature monitoring datasets at small scales, such as the long-term ecological research site at Lake Mendota, Wisconsin, and long-term low-frequency datasets such as the University of Rhode Island Watershed Watch program that samples 120 lakes across Rhode Island (Hollister et al. 2021). The introduction of the Water Quality Portal (WQP) aggregated surface water data, including temperature, across the United States (Read et al. 2017). However, 95.7% of the 275,897 National Hydrography Dataset (NHDPlus V2) sampled contiguous U.S. waterbodies in the WQP did not include in situ temperature measurements (Schaeffer et al. 2018b). Efforts to establish a national lake database have made progress but a comprehensive national scale database of lake surface water temperature is still lacking. In 2015, LAke multi-scaled GeOSpatial and temporal database of the Northeastern U.S. (LAGOS-NE), a multi-state lake database, was released (Soranno et al. 2017; Soranno et al. 2015). However, due to difficulties related to harmonizing the data across multiple sources and sampling methods, temperature was omitted from this database. A LAGOS-US database was later released for the contiguous U.S. with in situ lake temperature data sourced from the WQP up to 2021 (Cheruvelil et al. 2021; Smith et al. 2021).

Satellite remote sensing offers a method to increase both spatial and temporal coverage of lake temperature records, filling gaps inherent in in situ monitoring schemes. Several satellite platforms include thermal infrared bands capable of detecting surface temperature at various spatial and temporal resolutions. Since these data represent the thermal radiation leaving the water surface, the measurement is termed skin temperature, which tends to vary only slightly from bulk water temperature—generally by < 1 °C (Wilson et al. 2013). Many satellites provide daily coverage with relatively coarse spatial resolution, including the Advanced Very-High-Resolution Radiometer (AVHRR, 1.1 km) and the Moderate Resolution Imaging Spectroradiometer (MODIS, 1 km). Though their coarse resolution is better suited for marine applications, these platforms have been used to monitor surface water temperature of larger lakes (Bolgrien and Brooks 1992; Crosman and Horel 2009). Other satellites with finer spatial resolution are more commonly used for monitoring surface water temperature of inland lakes. The Advanced Spaceborne Thermal Emission and Reflection Radiometer (ASTER) has enabled lake surface temperature retrieval (Becker and Daw 2005) with 90-m spatial resolution on a 16-day basis since the launch of Terra in 1999. Terra will cease operation in 2027, having already exceeded mission length expectations. ECOsystem Spaceborne Thermal Radiometer Experiment on Space Station (ECOSTRESS) provides 70-m spatial resolution on a tasked basis, with revisit frequency ranging from hours to 5 days (Fisher et al. 2020) and is an experimental sensor. Launched in 2018, it lacks the robust historical record offered by other platforms such as Landsat. The Landsat series has included one or more thermal bands ranging in spatial resolution from 60 to 120 m since 1982. Currently, Landsat 8 and 9 thermal infrared sensors are delivering 100-m thermal data oversampled to a 30 m grid with a combined repeat of 8 days (Masek et al. 2020).

Schaeffer et al. (2018b) reported validation of an operational satellite provisional surface water temperature from the Landsat Analysis Ready Data. Open water pixels performed well, indicating Landsat data may supplement traditional in situ data by providing measures for most U.S. lakes, reservoirs, and estuaries (Clark et al. 2017; Schaeffer and Myer 2020) over consistent seasonal intervals for an extended period of record. Following the validation of Landsat surface water temperature, Myer et al. (2020) applied these data to a cyanoHAB forecast model across Florida lakes reporting 82% overall accuracy in the cyanoHAB prediction dataset. The Myer et al. (2020) cyanoHAB forecast model was applied to all Florida lakes that were resolved with the 300-m pixel resolution of the Sentinel-3 Ocean and Land Colour Instrument (OLCI) sensor (Donlon et al. 2012), which is used to generate satellite-derived measures of cyanobacteria that are publicly accessible through the Cyanobacteria Assessment Network (CyAN; Schaeffer et al. 2015; Seegers et al. 2021). However, Myer et al. (2020) had to supplement weekly gaps in the Landsat surface water temperature data using monthly climatology due to cloud cover and non-uniform scene acquisition across Florida. There are also known caveats to Landsat—specifically, inland waters may include missing data due to gaps in the Advanced Spaceborne Thermal Emission and Reflection Radiometer Global Emissivity Database. Additionally, errors have been reported in the surface temperature retrievals due to clouds and cloud shadows, according to one study investigating the Great Lakes (Cook et al. 2014). Prior to expanding the Myer et al. (2020) cyanoHAB forecast model beyond Florida lakes to forecast all 2,192 OLCI resolved lakes, a solution was required to fill in the spatial and temporal gaps in surface water temperature as a critical predictor.

Modeling can provide a useful method to estimate surface water temperature across unmeasured locations and fill temporal gaps in measured locations, where observations are often sparse. Increasing the spatial representation of lakes and the temporal continuity of temperature data would benefit efforts to forecast water quality events such as cyanoHABs. Temperature prediction models may be built using water temperature measures from either 1) traditional in situ measurements or 2) satellite data to achieve broader spatial representation and temporal continuity. In situ measures are more accurate but are limited in both temporal and spatial coverage. In contrast, Landsat satellite data provide measures of surface water temperature for all lakes, seasons, and temperature ranges, but with generally higher error than in situ measures.

Here, we demonstrate random forest models to provide a daily predicted surface water temperature dataset for all OLCI resolved lake locations. Random forest models are the state–of–the–art choice for tabular environmental data because they perform competitively with, or better than, deep neural networks (Borisov et al. 2024). Deep learning models are frequently challenged on real–world tabular data that are often noisy, complex and irregular, and performance depends on preprocessing, whereas random forest tree–based methods tolerate missing values, outliers, and capture heterogeneous feature importance. Meta–analyses and broad benchmarks corroborate this pattern, where tree–based methods outperform non–tree models across diverse domains and remain state–of–the–art on medium–sized tabular datasets while being computationally efficient (Grinsztajn et al. 2022; Uddin and Lu 2024).

The objectives of this study were to (1) determine the range and temporal and spatial representation of available Landsat and in situ temperature measures available for to the 2,192 OLCI resolved lakes, (2) validate Landsat surface water temperature performance based on concurrent reference in situ data and cloud presence, and (3) build and assess separate random forest models trained with in situ or Landsat surface water temperature data to compare the abilities of these approaches to provide spatiotemporally synoptic temperature records. A successful temperature model would provide comprehensive spatial and temporal temperature dynamics of each lake, for each week of the year, that supports national cyanoHAB forecasting. This study represents a novel contribution to temperature forecasting because it (1) validates Landsat surface water temperature with the presence of clouds, and (2) expands existing surface water temperature data coverage to match weekly measures spatially and temporally for OLCI resolved lakes supporting cyanoHAB forecasting.

## Methods

This section overviews the study methods and workflow (Fig. 1), including Landsat and in situ measured surface water temperature, satellite validation, random forest models, and input variables. All work was performed using RStudio version 4.4.1 and all spatial data were in the Albers 1983 projection. Code used for the analysis, including cached model runs, are available from https://github.com/usepa/sw_model.

**Figure 1.**
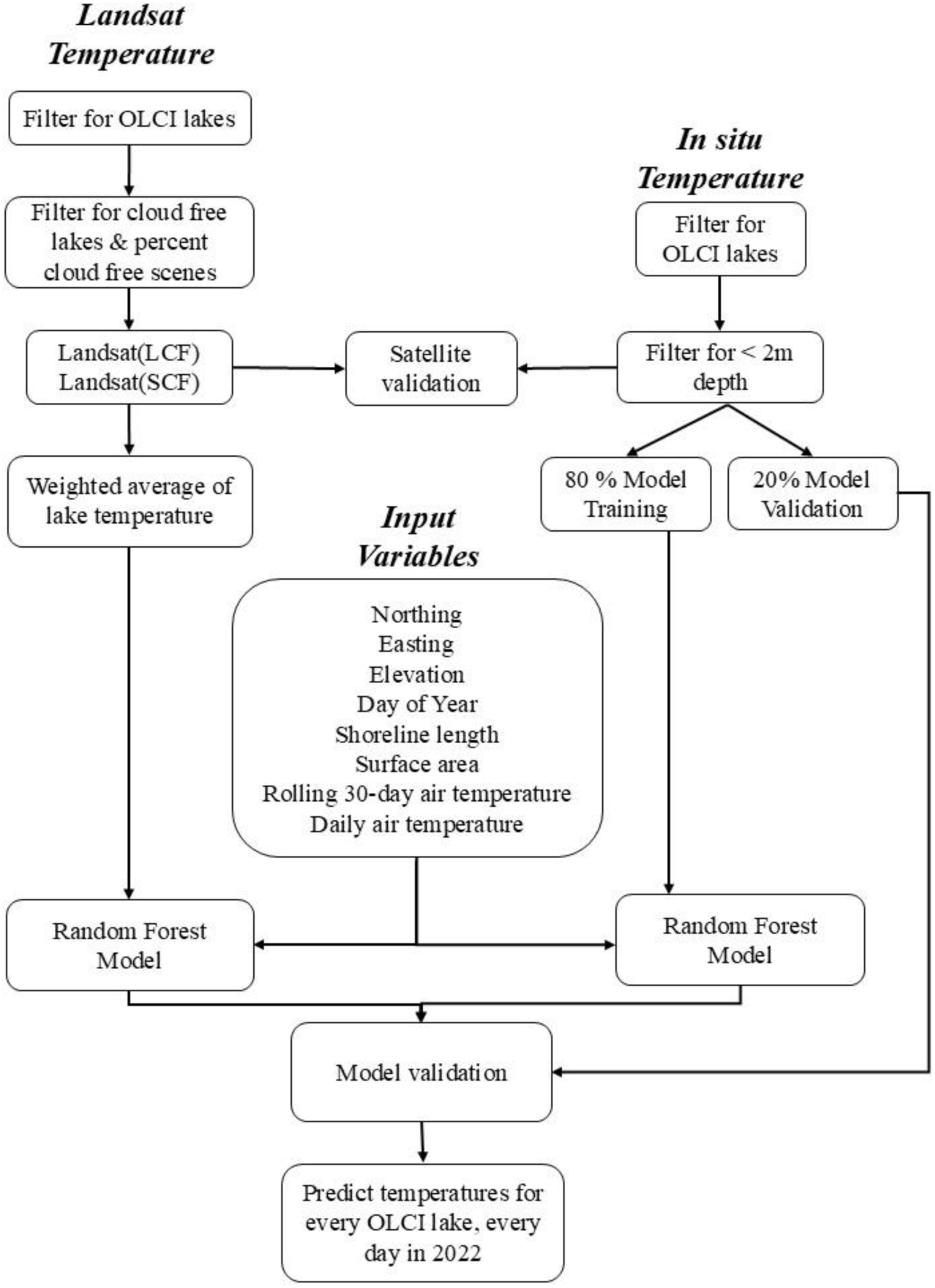
Schematic workflow diagram for analysis including Landsat and in situ surface water temperature, satellite validation, input variables, and random forest models corresponding to each subsection in the methods.

### Sentinel-3 OLCI satellite resolved lakes

Sentinel-3 OLCI resolved lakes and reservoirs were previously identified by Clark et al. (2017) based on the National Hydrography Dataset Plus version 2.0 database (McKay et al. 2012), where the categorization of lakes included coastal lagoon systems. Briefly, resolved lakes were identified based on the minimum Euclidean distance from shore that will accommodate a 300 m satellite pixel. This approach was updated with snow and ice masking using the Iterative Multisensor Snow and Ice Mapping System Northern Hemisphere Snow and Ice Analysis data from the National Snow and Ice Data Center as described in Urquhart and Schaeffer (2020). Briefly, all satellite pixels within the spatial area of the snow/ice mask were omitted from further lake water quality analysis. A minimum of three 300 m valid water pixels were considered OLCI resolved lakes (Schaeffer et al. 2022). The period for this study was 2007, 2012, and 2016-2022 which coincided with available in situ water temperature observations.

### In situ temperature

In situ discrete measurements of surface water temperature were accessed through the National Water Information System (NWIS, www.waterqualitydata.us) and the National Lakes Assessment (NLA) (U.S. EPA 2009; U.S. EPA 2011) because they were consistently high quality and collected with uniform methods required for temperature models (Kilpatrick et al. 2001). NWIS data were based on the following criteria: lake, reservoir, or impoundment site type; water sample medium; physical characteristic group; and water temperature characteristic. NWIS data used were from the period of January 1, 2016 through December 31, 2022 within the contiguous United States (CONUS). NLA data used were from the summer months of 2007, 2012, and 2017. Discrete samples were retained if they were within an OLCI resolved lake and in the upper 2 m of the water column, representing the modeled target of a mean daily temperature and the typical red and near-infrared satellite spectral band penetration depth of the cyanobacteria monitoring (Mishra et al. 2005; Wynne et al. 2010). The maximum number of samples per lake was limited to 250 observations to prevent a model training bias with a small subset of lakes that were dominant with in situ samples. The observations were randomly selected for lakes with more than 250 total samples. These remaining in situ data were randomly split with 80% applied in model training and 20% used for model validation. Any in situ data coincident with the same calendar day Landsat overpass were applied to validate the satellite surface water temperature data to maximize the potential number of match-ups.

### Landsat temperature

Landsat Surface Temperature data were obtained directly from the United States Geological Survey (USGS) Analysis Ready Data (ARD) repository (Cook et al. 2014; Hulley et al. 2014; Hulley and Hook 2009; Hulley et al. 2015). Landsat ARD provided uniform and systematic observation data across multiple satellite missions in a structured grid output for Landsat 7, 8, and 9 as well as quality assurance information from the study period of May 2016 to April 2022. Landsat ARD scenes were accessed for the entire conterminous United States using the USGS Machine-to-Machine API. To ensure completeness of the dataset, downloaded Landsat files were compared to the USGS bulk metadata file (https://www.usgs.gov/landsat-missions/bulk-metadata-service) for the study period. Landsat data were first filtered to the OLCI resolved lakes. Landsat surface water temperature was masked using the QA_PIXEL band to only retain pixels with QA bits categorized as “clear”, “low cloud”, and “low shadow”. Previous efforts report that Landsat data result in errors as large as several degrees C during an unstable atmosphere and cloudy conditions (Laraby et al. 2016). Therefore, any observations containing clouds or cloud shadow flags within the lake were discarded from further analysis. Lakes that were cloud and cloud shadow free were retained for analysis even if scenes contained these flags but were outside the lake area. An additional mask was applied to pixels within 300 m (equivalent of one 300 m Sentinel-3 OLCI pixel) from the NHD defined static shoreline to avoid pixels contaminated by a land signal for the Landsat Thematic Mapper, Enhanced Thematic Mapper Plus, and Thermal Infrared Sensors (Schaeffer et al. 2018b; Urquhart and Schaeffer 2020). Pixels below 0°C were also removed from further analysis to avoid frozen waters. The 1 km North America mask from the U.S. National Ice Center’s Interactive Multisensor Snow and Ice Mapping System was used to mask for snow and ice presence (National Ice Center 2023; Urquhart and Schaeffer 2020). On a given day, lakes identified as having snow or ice from the National Ice Center’s mask were removed from further consideration. Remaining non-masked lake surface temperature pixels were used for analysis with a single weighted average for each lake on each day an image was available. The cloud free lakes are henceforth labelled Landsat_(LakeCloudFree)_.

Despite using the QA band provided by USGS to remove clouds and cloud shadows within lakes, Landsat can have cloud interference across the scene as previously demonstrated in coastal oceans and the Great Lakes (Cook et al. 2014; Laraby and Schott 2018). Cloud interference was not considered in an initial analysis of lake water temperature that used scenes relatively free of cloud contamination (Schaeffer et al. 2018b). Therefore, we tested if errors propagated across a scene from cloud and cloud shadow. A second filtering was applied to Landsat_(LakeCloudFree)_ scenes with cloud or cloud shadow presence using <25%, <10%, and <1% cloud cover and shadow. The resultant <1% cloud and cloud shadow scenes are henceforth labelled Landsat_(SceneCloudFree)_.

### Satellite validation

In situ and Landsat comparisons represented a diverse geographic area under a variety of atmospheric and aquatic conditions (Kilpatrick et al. 2001) to ensure a comprehensive validation dataset. A temporal restriction of same-calendar-day matchups between Landsat overpass and in situ data were used to maximize the number of potential in situ measures matched with satellite observations while minimizing the complexities of bio-physical changes such as vertical and horizontal movement of the water column. In situ measures were matched to a single 30 m oversampled pixel (1 × 1 pixel array) at the discrete in situ sample location. The validation was applied to all Landsat cloud free lakes with scenes regardless of cloud cover shadow, and with scenes containing <25%, <10%, and <1% cloud cover and shadow. Performance statistics included the mean absolute error (MAE), mean absolute percentage error (MAPE), and bias.

### Input variables

Random forest model input variables were northing and easting of the lake centroid, elevation, day of year, static shoreline length, static surface area, rolling 30-day air temperature prior to the prediction date, and daily air temperature on the date of the prediction. Lake centroid northing and easting were obtained from the National Hydrography Dataset Plus version 2.0 database (McKay et al. 2012). Lake elevation was obtained from the Lake-Catchment dataset (Hill et al. 2018). Lake surface area and lake shoreline length were calculated using the lakemorpho R package (Hollister and Stachelek 2017; https://github.com/jhollist/lakemorpho). Parameter-elevation Regressions on Independent Slopes Model (PRISM) 4 km air temperature data for North America were downloaded from the Oregon State PRISM Climate Group (PRISM 2022; https://prism.oregonstate.edu/) for 2007, 2012, and 2016-2022. Daily and averaged 30-day prior air temperature were obtained from the PRISM data for each OLCI resolved lake.

### Random Forest Model

Random forest models were trained on in situ surface water temperature and Landsat surface temperature for all OLCI resolved lakes using the *randomForest* R package (Breiman 2001; Liaw and Wiener 2002). Surface water temperature was modeled with three random forest models and 100 trees each using (1) the Landsat_(LakeCloudFree)_, (2) the Landsat_(SceneCloudFree)_, and (3) in situ surface water temperature measures. The in situ dataset was randomly split into 80% training and 20% validation. The *randomForest* function calculated the coefficient of determination (R^2^) and mean-square error that was converted to root-mean-square error (RMSE). The *randomForest* predicted output was used to determine the out-of-bag predictions for bias and mean absolute error (MAE). MAE and bias (MAE_applied_ and bias_applied_) were calculated for the validation data. Each of the three models were used to predict daily averaged water temperature for all OLCI resolved lakes in 2022 as a demonstration and to test for expected seasonal trends as a quality check.

## Results

### Satellite validation

Landsat surface water temperature was validated with in situ surface (< 2 m) water temperatures by matching satellite single 30 m oversampled pixel values with same calendar day coincident in situ samples. These matchups were evaluated with different amounts of cloud cover and cloud shadow in the Landsat scene. All the in situ data matched with Landsat_(LakeCloudFree)_ had 563 matchups with an MAE of 2.99°C, MAPE of 14%, and bias of -1.3°C (Fig. 2a). When cloud cover and cloud shadow were reduced to < 25% in the Landsat scenes, 388 matchups remained, with improved MAE of 2.07°C, MAPE of 11%, and bias of -0.31°C (Fig. 2b). Reduced cloud cover and shadow of < 10% improved error, resulting in 296 matchups with MAE of 1.87°C, MAPE of 10%, and bias of -0.2°C (Fig. 2c). Landsat_(SceneCloudFree)_ was defined as < 1% cloud cover and shadow, and resulted in 149 matchups and MAE of 0.94°C, MAPE of 6%, and bias of +0.39°C (Fig. 2d). The improvement in validation metrics with the reduction of Landsat scene cloud cover and shadow indicated that the Landsat_(SceneCloudFree)_ would provide the lowest error with a small positive bias.

**Figure 2.**
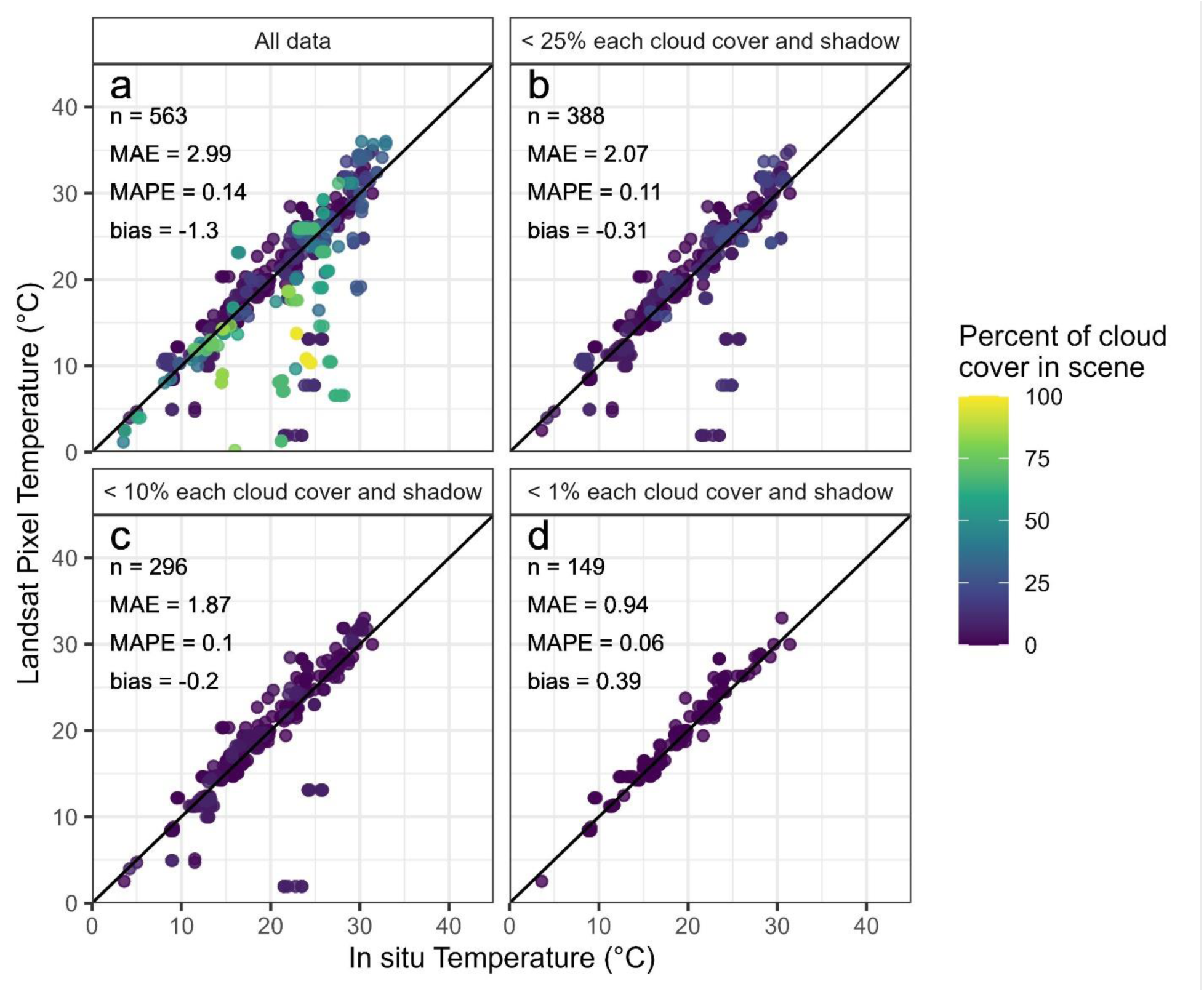
Validation scatterplots for cloud free lakes with (a) scenes regardless of cloud cover and shadow, (b) scenes containing < 25 %, (c) < 10 %, and (d) < 1 % cloud cover and shadow. Matchups were single pixel and same calendar day with in situ measures where n is the number of validation points. There were no available matchups with 0% cloud and cloud shadow scenes.

### Model training and validation

The total number of in situ observations was 15,754 across 156 of the 2,192 OLCI resolved lakes. The maximum number of samples per lake was limited to 250 observations to prevent model training bias. Randomly removed observations for lakes with more than 250 observations decreased the number of in situ observations to 8,645. The in situ training dataset contained 6,743 samples from 156 lakes (Fig. 3a) and the validation set had 1,636 samples from 126 lakes (Fig. 3b). Landsat_(LakeCloudFree)_ had the most observations across more lakes than Landsat_(SceneCloudFree)_ (Fig. 3), with 298,401 observations across all 2,192 OLCI resolved lakes (Fig. 3c). Landsat_(SceneCloudFree)_ filtering decreased the number of observations to 270 across 126 OLCI resolved lakes (Fig. 3d). With the exception of Maine, few Landsat_(SceneCloudFree)_ data came from the eastern CONUS. In situ training data ranged from 1 to 217 observations averaged per lake, Landsat_(LakeCloudFree)_ had 12 to 417 observations averaged in time per lake to estimate the mean daily in situ water temperature, Landsat_(SceneCloudFree)_ ranged from 1 to 26 observations, and the in situ validation range was between 1 and 20 observations per lake.

**Figure 3.**
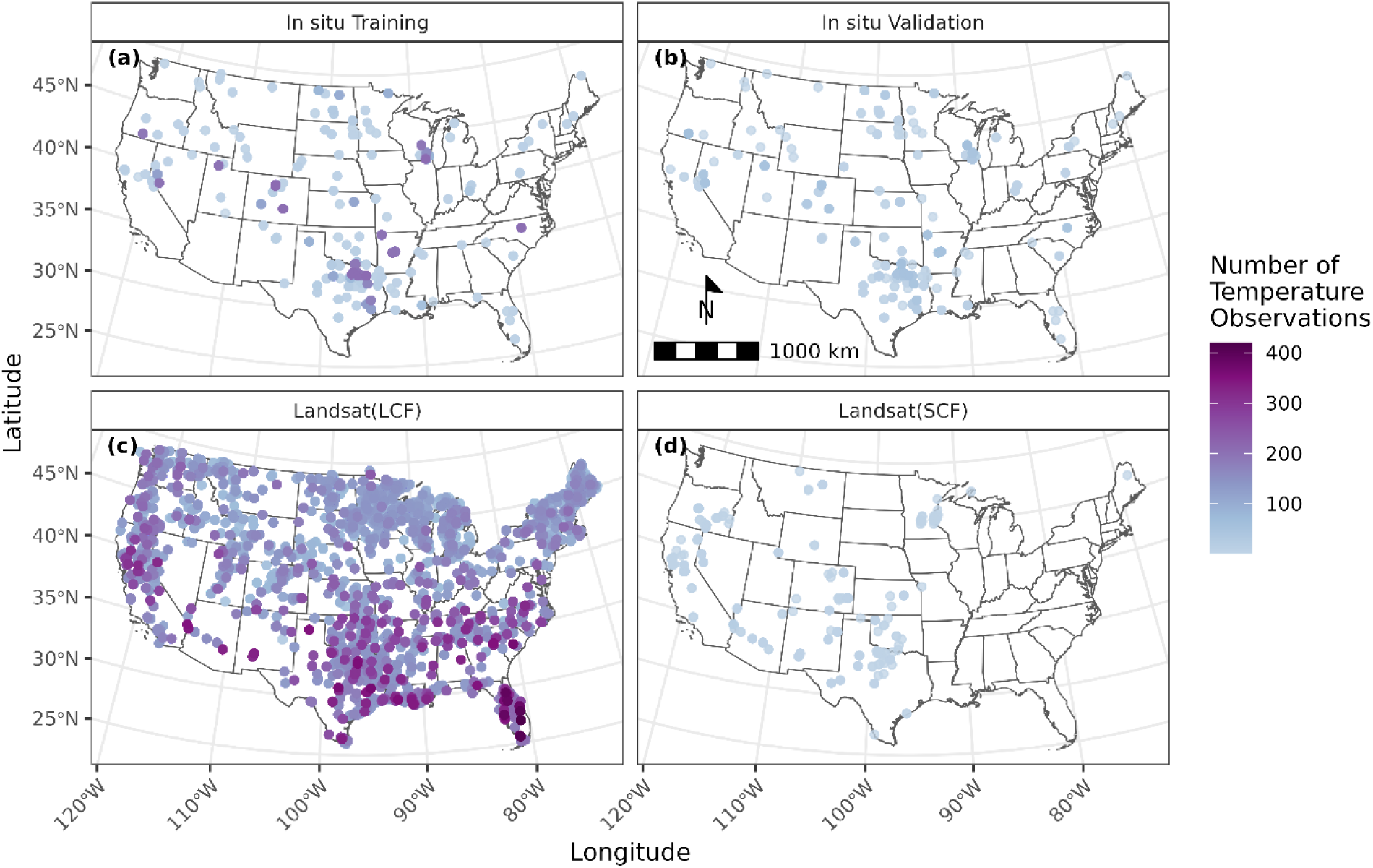
Number of temperature observations per lake for (a) in situ training, (b) in situ validation, (c) Landsat_(LakeCloudFree)_, and (d) Landsat_(SceneCloudFree)_datasets. Landsat_(LakeCloudFree)_ and Landsat_(SceneCloudFree)_ are shown as Landsat(LCF) and Landsat(SCF), respectively.

Since the in situ, Landsat_(LakeCloudFree)_, and Landsat_(SceneCloudFree)_ datasets had differing numbers of temperature observations (Table 1), the corresponding predictor variables also have different ranges (Fig. 4). All four of the datasets had the most observations during warmer periods with medians from 17.8°C to 22.8°C (Fig. 4a and 4b). Temperature distribution was higher for the Landsat_(SceneCloudFree)_, whereas in situ and Landsat_(LakeCloudFree)_ temperatures and distributions were similar. Landsat_(SceneCloudFree)_ was skewed toward the west with a median of – 103.79 degrees longitude (easting: -667416 meters) compared to the other datasets (Fig. 4c). Landsat_(LakeCloudFree)_ had the broadest distribution across latitudes with a higher median of 42.51 degrees (northing: 2171446 meters). All four datasets provided representation across all months of the calendar year (Fig. 4e). The distribution of day of year frequencies was similar for all datasets, with the highest frequencies occurring between day 150 and 250. Elevations ranged from < 10 m to sub-alpine ranges of 1,500 m to 3,600 m; Landsat_(SceneCloudFree)_ was higher in elevation with a median elevation at 1089 m compared to the other three datasets (Fig. 4f). Lake area (Fig. 4g) and shoreline length (Fig. 4h) were similar with median area of 10.1 and 9.6 km^2^ and median shoreline length of 28.5 and 32.7 km for the Landsat datasets. The in situ datasets were higher with median area of 53.0 km^2^ and median shoreline length of 127.7 km (Fig. 4g and 4h).

**Figure 4.**
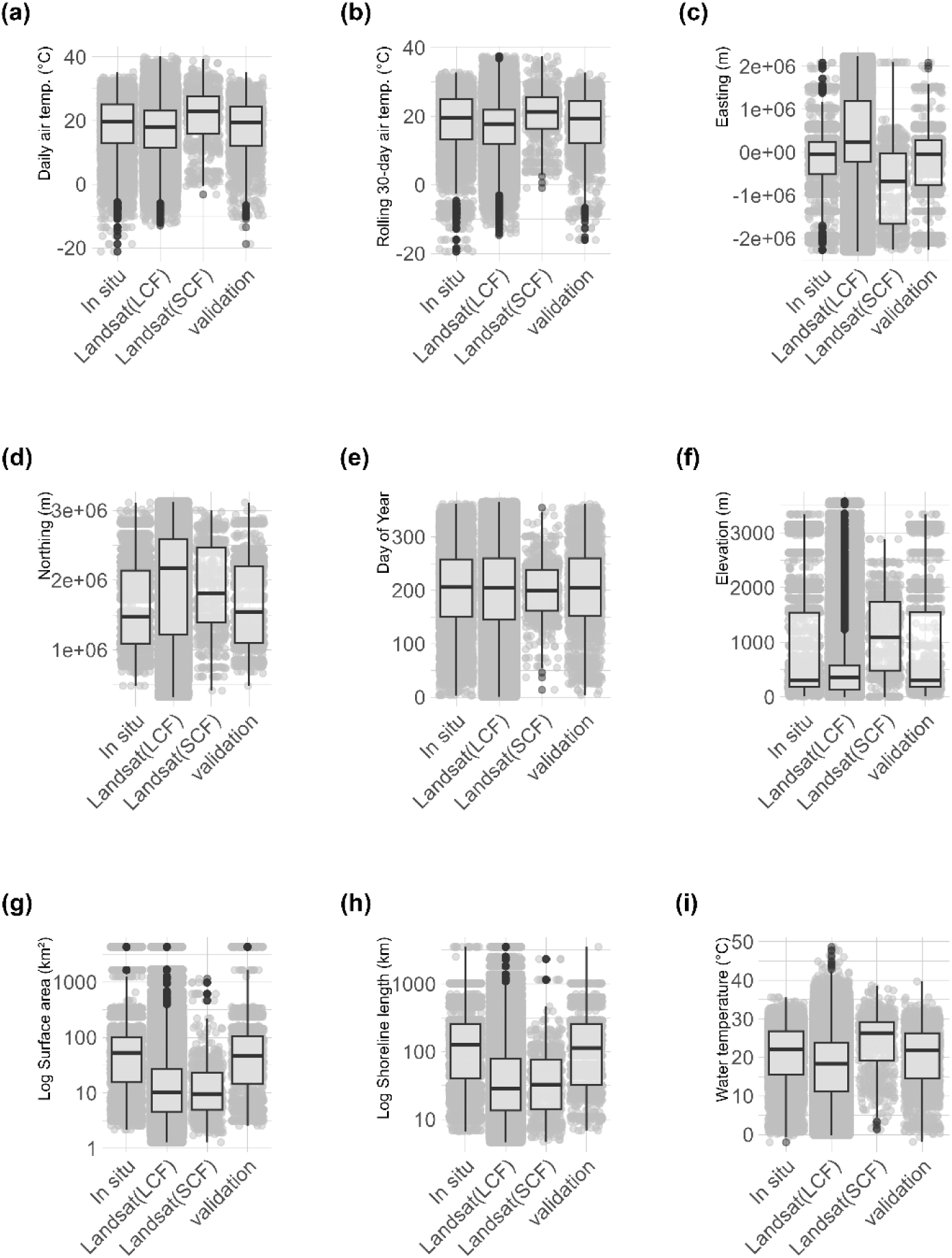
Distributions of (a) average temperature, (b) prior 30-day average temperature, (c) easting projected coordinates, (d) northing projected coordinates, (e) day of year, (f) elevation, (g) lake area, (h) lake shoreline length, and (i) water temperature for Landsat_(LakeCloudFree)_, Landsat_(SceneCloudFree)_, in situ training, and in situ validation data. Grey points in background are the individual observations with a small randomization factor added to reduce over plotting. Landsat_(LakeCloudFree)_ and Landsat_(SceneCloudFree)_ are shown as Landsat(LCF) and Landsat(SCF), respectively. Box range was 25^th^ and 75^th^ percentiles, line was the median, and black points were outliers.

**Table 1.** Performance metrics for the Landsat_(LakeCloudFree)_, Landsat_(SceneCloudFree)_, and in situ models for random forest out of bag observations.

| Metric | Model Variable |  |  |
| --- | --- | --- | --- |
|  | Landsat<sub>(LakeCloudFree)</sub> | Landsat<sub>(SceneCloudFree)</sub> | in situ |
| R<sup>2</sup> | 0.71 | 0.71 | 0.96 |
| RMSE (°C) | 4.43 | 4.22 | 1.51 |
| Bias (°C) | -0.07 | 0.07 | 0.04 |
| MAE (°C) | 2.98 | 2.48 | 1.02 |
| n | 298,401 | 270 | 1,907 |

### Model Output

All three models had similar responses over the seasons of 2022 (Fig. 5). Minimum temperatures were predicted for winter days between January (day 0) through mid-February (day 49) and in December. Temperatures increased from the end of February (day 50) and peaked in July (day 200). From day 200 through December temperature declined across all models. The Landsat_(LakeCloudFree)_ model regularly predicted cooler temperatures across the entire 2022 year compared to the in situ and Landsat_(SceneCloudFree)_ models. Landsat_(SceneCloudFree)_ generally predicted higher temperatures throughout the 2022 year than the in situ and Landsat_(LakeCloudFree)_ models, particularly in the cooler months. The closest agreement between the Landsat_(SceneCloudFree)_ and in situ models occurred during the shoulder seasons from May days 125 through 150 and August to October days 225 through 300. Landsat_(SceneCloudFree)_ had the greatest difference from in situ in the winter months from days 0 through 125 and again after day 300.

**Figure 5.**
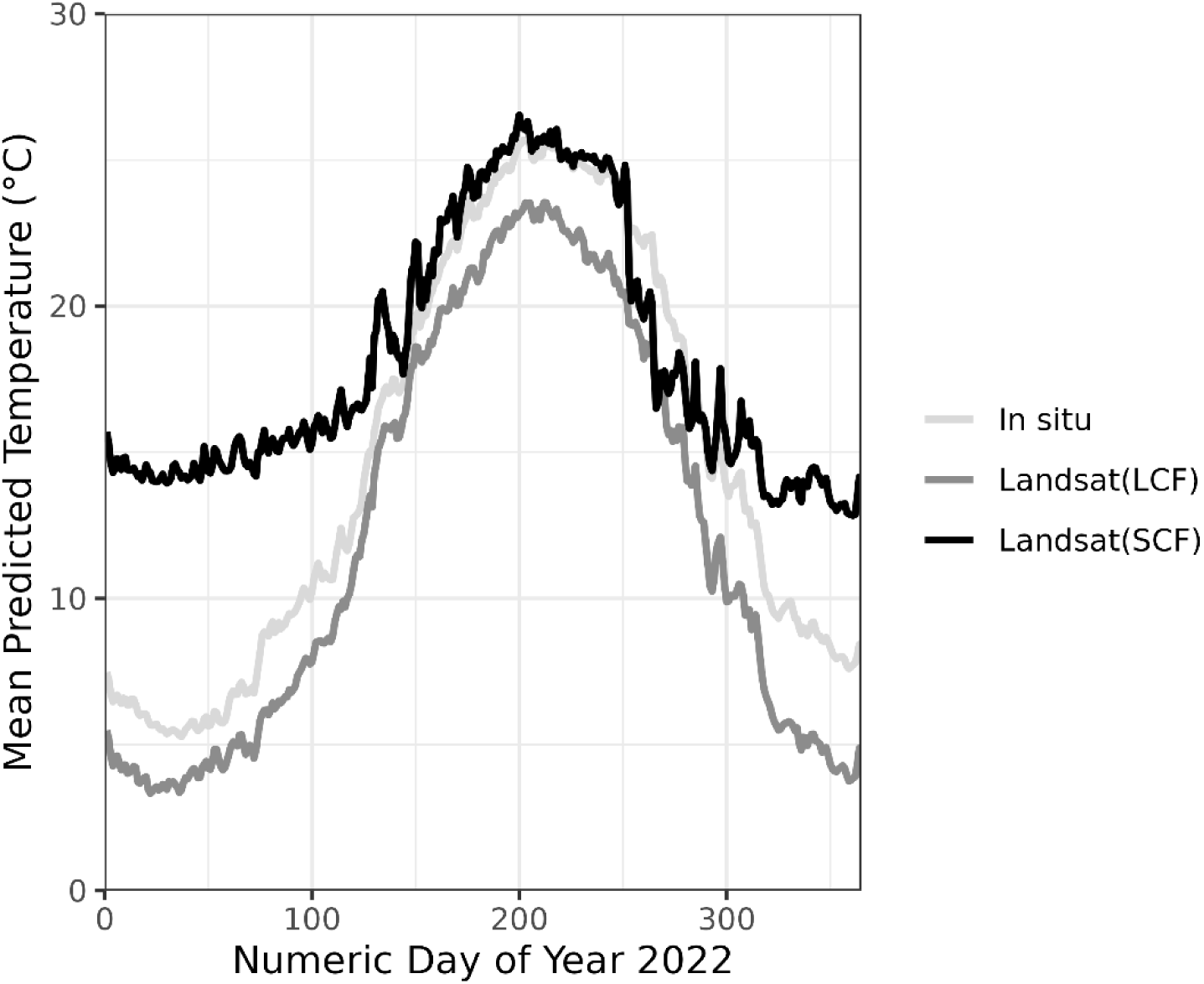
Demonstration of mean daily predicted water temperature for all OLCI resolved lakes in 2022 from the Landsat_(LakeCloudFree)_, Landsat_(SceneCloudFree)_, and in situ models. Landsat_(LakeCloudFree)_ and Landsat_(SceneCloudFree)_ are shown as Landsat(LCF) and Landsat(SCF), respectively.

Easting, elevation, and lake surface area were the three most important variables for the Landsat_(LakeCloudFree)_ random forest model (Fig. 6a). The daily average air temperature and Northing were in the top three most important variables for the Landsat_(SceneCloudFree)_ (Fig. 6b) and in situ (Fig. 6c) models. The variable importance for the in situ model had rolling 30-day air temperature and daily average air temperature along with day of year as the most important variables.

**Figure 6.**
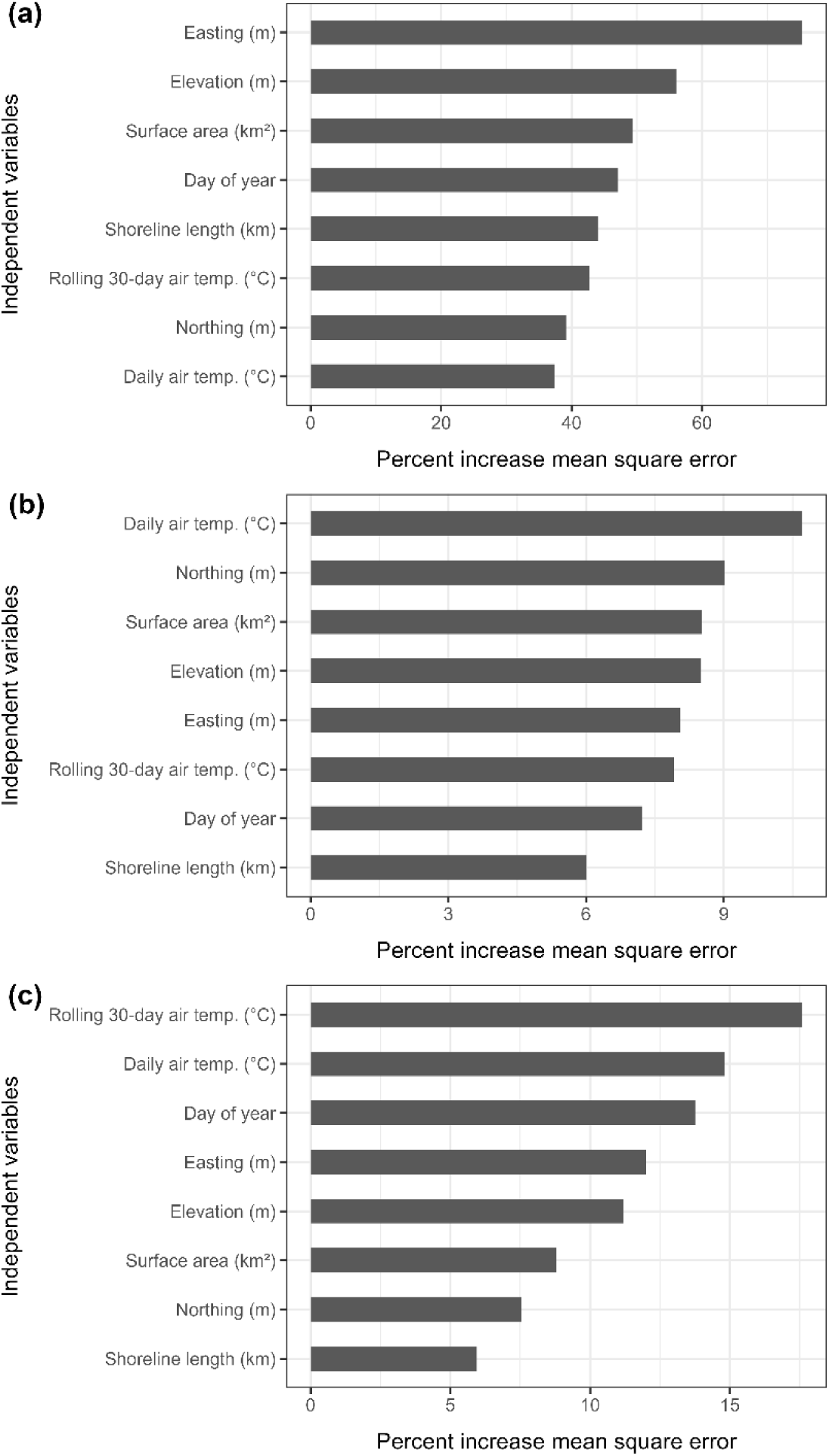
Variable importance plots for the (a) Landsat_(LakeCloudFree)_, (b) Landsat_(SceneCloudFree)_, and (c) in situ random forest models.

The partial dependency plots show the marginal effect of each variable on the predicted temperature (Fig. 7). Consistent with other results (i.e. Fig. 5), the Landsat_(SceneCloudFree)_ model tended to have the warmest predictions and the Landsat_(LakeCloudFree)_ model had the coolest predictions. The marginal effect is similar across the three models for the average air temperature (Fig. 7a), 30-day prior average air temperature (Fig. 7b), and day of year (Fig. 7c). Warmer air temperatures result in warmer predicted lake temperature and these level off at temperatures above ∼20-25 °C for both average air temperature variables. Day of year follows the expected warming through the course of the year with warmest temperatures in summer and coolest in winter. The east-west change in predicted temperature is generally less than 1 °C across the full range of easting (Fig. 7 d), except for Landsat_(SceneCloudFree)_. Predicted temperature decreases at higher latitudes for in situ and Landsat_(LakeCloudFree)_ (Fig. 7e), while Landsat_(SceneCloudFree)_ shows an increase at lower latitudes and then is mostly invariant. The two lake morphometry metrics behave similarly (Fig. 7f and 7g). The Landsat_(LakeCloudFree)_ and Landsat_(SceneCloudFree)_ models show slightly warmer predicted temperatures in smaller lakes with a larger impact on lake size shown with the Landsat_(SceneCloudFree)_ model. The in situ model shows little variation across most of the range of lake sizes. It is important to note that most of the samples were concentrated in smaller lakes; lakes with area greater than 1000 km^2^ and 1000 km or more of shoreline were rare. The effect of elevation on predicted lake temperatures is mostly similar across models, with slightly lower predicted temperatures at higher elevations, but both Landsat_(SceneCloudFree)_ and in situ show an abrupt decrease at 2000 meters and 3000 meters, respectively (Fig. 7h).

**Figure 7.**
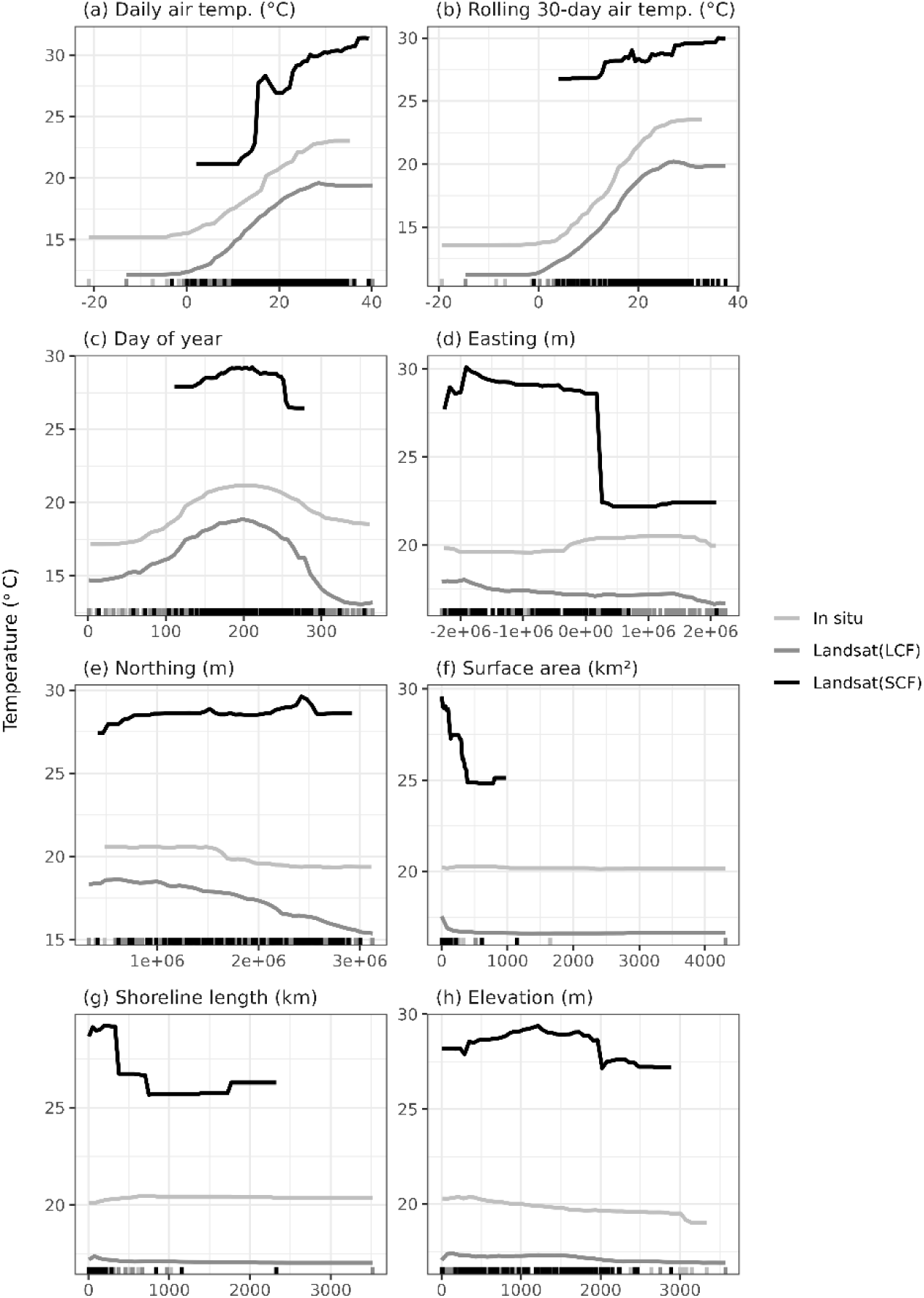
Partial dependency plots for the Landsat_(LakeCloudFree)_, Landsat_(SceneCloudFree)_, and in situ random forest models. The tick marks along the x-axis show the distribution of the training data values from the 0^th^ to 100^th^ percentiles.

### Model Performance

The *randomForest* predicted output was used to determine the out-of-bag R^2^, RMSE, bias and MAE (Table 1). After applying the models to the validation dataset we calculated MAE_applied_ and bias_applied_ to quantify performance (Fig. 8). While error metrics generated from out-of-bag observations are in theory independent since they are based on observations held out from each individual tree, the validation data were not used in any trees and were thus completely independent. Therefore, the applied validation metrics likely provide a more accurate assessment of model performance (Janitza and Hornung 2018; Millard and Richardson 2015). The in situ model had the lowest MAE_applied_ followed by Landsat_(LakeCloudFree)_ and Landsat_(SceneCloudFree)_ models. Landsat_(SceneCloudFree)_ had a small positive validation bias_applied_ of 0.81 °C, while Landsat_(LakeCloudFree)_ had a negative bias_applied_ of -1.97 °C. Error metrics indicated that the in situ model had the best fit and best overall performance.

**Figure 8.**
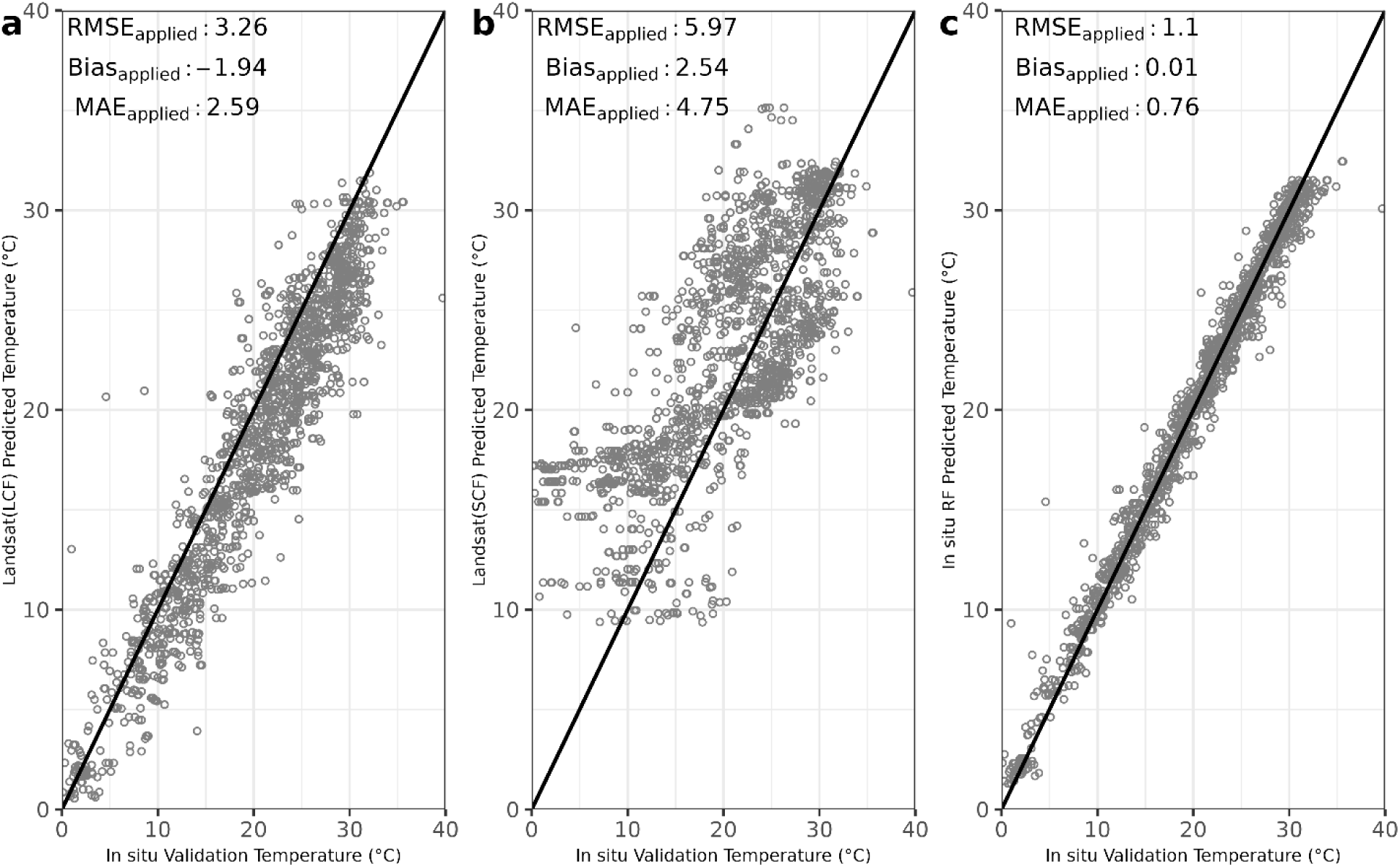
Predicted temperatures for (a) Landsat_(LakeCloudFree)_, (b) Landsat_(SceneCloudFree)_, and (c) in situ models compared to the in situ validation dataset, with 1:1 lines shown. Landsat_(LakeCloudFree)_ and Landsat_(SceneCloudFree)_ are shown as Landsat(LCF) and Landsat(SCF), respectively. There were 1,636 (N) validation points.

The majority of waterbody average error was > 3 °C in both Landsat models (Fig. 9). The Landsat_(LakeCloudFree)_ model showed nearly uniform cooler predicted temperatures across the U.S. The Landsat_(SceneCloudFree)_ model overestimated temperature by > 3 °C across much of the western half of the country, particularly in higher latitude lakes. Landsat_(SceneCloudFree)_ errors in the eastern half of the country tended to underestimate temperature by > 3 °C. Regions of negative error from the Landsat_(SceneCloudFree)_ model appear to coincide with data poor locations (Fig 3b). The in situ model had far fewer lakes with error > 3 °C and no discernable cold or warm bias spatially.

**Figure 9.**
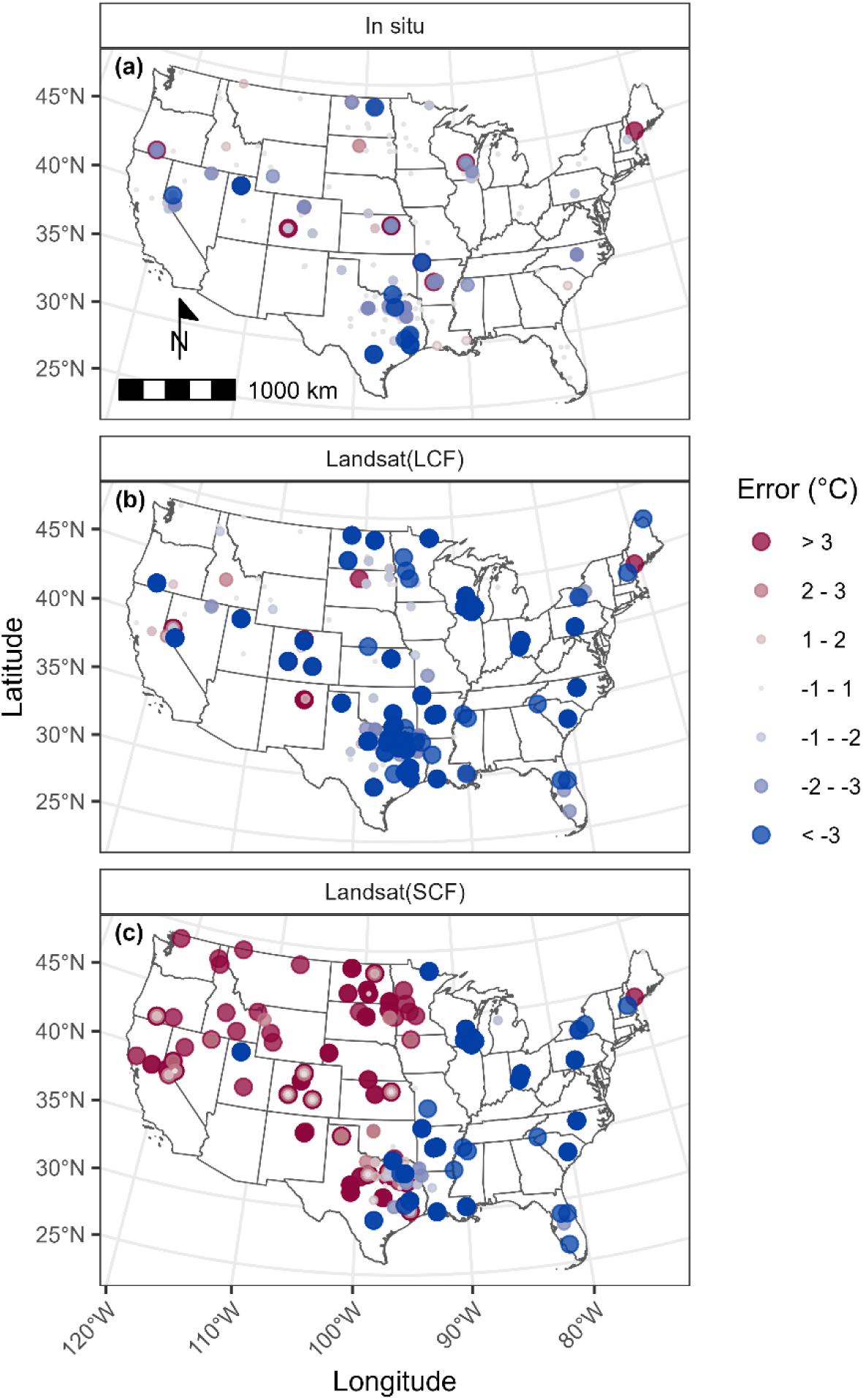
Average error per lake when compared to the in situ validation dataset for the (a) in situ, (b) Landsat_(LakeCloudFree)_, and (c) Landsat_(SceneCloudFree)_ trained random forest predicted temperatures. Landsat_(LakeCloudFree)_ and Landsat_(SceneCloudFree)_ are shown as Landsat(LCF) and Landsat(SCF), respectively.

## Discussion

Our results align with previous studies highlighting the challenges of using satellite data for inland water temperature monitoring due to cloud cover, proper corrections, and algorithm implementations. Similar to Cook et al. (2014), we observed increased error and negative bias with more in scene cloud cover in the Landsat data based on comparisons with in situ measurements. Though the Landsat dataset offered far more observations than the in situ dataset, satellite temperature measurements contained several sources of error that may have impacted the quality of the product. Landsat data underwent atmospheric correction, a process that is known to introduce uncertainty, particularly over inland waters (Moses et al. 2017). It was evident that images containing any cloud cover—even in parts of the image distant from the area of interest—may contain elevated error, based on the improved performance of Landsat without filtering for cloud cover and shadow to <1 cloud cover and shadow. The cloud filtering performed here was done using the cloud flag included within Landsat, a product known to miss clouds, particularly thin clouds (Qiu et al. 2019). If such clouds were present in the Landsat_(SceneCloudFree)_ dataset, they could be responsible for some observed error, as well. Additionally, though observations near land were removed in this study, it was still possible that land adjacency could play a role in introducing error (Schaeffer et al. 2018b).

This study demonstrated the application of random forest models to predict lake surface water temperatures using both in situ and Landsat derived temperature data. We found better performance of the in situ model over the Landsat models, particularly the Landsat_(SceneCloudFree)_ model, which had higher MAE and bias. The Landsat temperature dataset offers a more comprehensive record with which to build models compared to the in situ dataset; however, the poorer performance of the Landsat_(LakeCloudFree)_ and Landsat_(SceneCloudFree)_ models suggests that error in the Landsat temperature dataset is high enough to compromise predictive power of models trained with it. Thus, our results confirm the importance of data quality in model construction. Comparing the two Landsat models, the Landsat_(SceneCloudFree)_ model showed higher error than the Landsat_(LakeCloudFree)_ model, despite being trained with more reliable data (Fig. 2). This may have been due to the smaller amount of training data (n = 149 vs. 563), and possibly poor representation across all seasons as a result (Fig. 4e). The dominance of higher elevations in Landsat_(SceneCloudFree)_ (Fig. 4f) may be attributed to a clear-sky sampling bias, as Mercury et al. (2012) showed that global mean cloud coverage from 2001 through 2010 exhibited orographic symmetry, with leeward mountain slopes experiencing systematically lower cloudiness than windward slopes. The drier regions of our study area also occurred at slightly higher elevations, and this cloud-related bias limited observations in lower elevations, such as along the coasts, constraining the spatial distribution of lakes. Consistent with this interpretation, Mercury et al. (2012) reported a westward decrease in cloud cover from the U.S. Great Plains to the leeward side of the Rocky Mountains, a pattern that closely matched the distribution of cloud-free Landsat scenes shown in our Figure 3d. This work validated Landsat surface water temperature under varying cloud conditions and developed a continuous predicted temperature dataset for all Sentinel-3 OLCI resolved lakes in CONUS. These advancements provide a comprehensive record of lake temperature, critical for predicting cyanoHABs.

Our study extends the findings of Kreakie et al. (2021) by incorporating a larger and more temporally diverse dataset, improving the model performance range below 15°C and improving the parameterization for all Sentinel-3 OLCI resolved lakes (Clark et al. 2017; Schaeffer et al. 2018a). By including NWIS and 2007, 2012, and 2017 NLA data, we were able to construct our in situ model with greater than four times the number samples used by Kreakie et al. (2021). The model temporal trends confirm the seasonal patterns reported in other studies (MacCallum and Merchant 2014; Piccolroaz et al. 2020). Warming typically occurred in early spring, between March through June, with a maximum from late June through beginning of September and declined late September through December (Fig. 5) Landsat_(SceneCloudFree)_ were warmer than Landsat_(LakeCloudFree)_ throughout the predicted 2022 calendar year, similar to results reported by Ermida et al. (2019; see their Fig. 10 for North America) due to clear-sky bias. Clear skies allow more solar radiation and therefore higher temperature values with positive bias.

Random forest models only perform well predicting values within the range of observed conditions, and the lack of wintertime data is a limit with the Landsat_(SceneCloudFree)_ model ability to predict cold conditions as demonstrated by elevated winter temperatures when compared to the in situ and Landsat_(LakeCloudFree)_ models. Comparing the in situ model to other similar approaches, we observed a slight improvement in performance. Using 2,282 in situ observations from the NLA, Kreakie et al. (2021) reported RMSE of 1.48°C, slightly higher than that reported here (1.11°C). Read et al. (2019) used Process Guided-Deep Learning models with temperature profile data to predict depth profiles with an RMSE of 1.65 °C. Earlier work that built a general linear model to predict maximum lake surface-water temperature for a collection of 872 Canadian lakes achieved a RMSE value of 2.53 °C (Sharma et al. 2007). Korver et al. (2024) validated Landsat-derived surface water temperature using lake center measures from 63 sites and 2,074 observations with RMSE of 1.71 °C. The Korver et al. (2024) dataset reported better performance with the Landsat models than either our Landsat_(LakeCloudFree)_ or Landsat_(SceneCloudFree)_ models, but our in situ model performed the best overall based on RMSE. The lower Landsat RMSE from Korver et al. (2024) may be a result of limiting the Landsat data extraction to the 50 m buffer at the center of each lake. Willard et al. (2022) reported long short-term memory deep learning model RMSE of 1.61 for 185,549 lakes in the United States. Tong et al. (2023) reported an MAE of 1.2 between their FLake model and satellite observations for 92,245 global lakes, whereas our Landsat models did not perform as well, but our in situ model MAE was lower at 0.76.

Our models, as with other temperature models, have 1.11-3.40 °C RMSE and this may limit the application depending on the science question. As an example, accuracy requirements for sea surface temperature to support climate monitoring and weather forecasting generally require uncertainty of 0.3 to 0.6 °C with long term stability (Donlon et al. 2007; Wick et al. 2002). Longer term climate impacts, such as the length of warmer seasons, long term increases of water temperatures, and warmer winter periods were not considered and were beyond the scope of this study as these results were reported previously (Arndt and Blunden 2017; Layden et al. 2015; Rose et al. 2016; Winslow et al. 2017; Woolway and Merchant 2017).

Easting and northing function as important variables (Fig. 6) because they proxy large–scale gradients that influence lake surface temperature. Easting may represent proximity to oceanic heat sources, atmospheric circulation, and climate zone spatial classification (Beck et al. 2018; Karl and Koss 1984). Northing reflects seasonality and temperature–density relationships controlling stratification (Kraemer et al. 2015) as well as climate zone spatial classification. Lake surface area impacts wind fetch and water residence time. Shoreline length impacts depth that waves can mix water and changes to water column stratification (Moses et al. 2011). Together, these variables serve as proxies for climatic variance and local physical controls not fully captured by radiative inputs alone, explaining their high importance in the temperature model.

The variable importance plots for both in situ and Landsat_(SceneCloudFree)_ are fairly similar (Fig. 7). Both the 30-day temperature average and daily temperature are in the top three important variables which we assume explains the impact of both seasonal warming trends and more immediate ambient temperature on lake temperature. Yet for the Landsat_(LakeCloudFree)_ model we see very different important variables selected with easting as the top importance variable. In Figure 7, the predicted temperature for the in situ model increased as the easting increased, which was not the pattern of the other two models. It is difficult to determine what exactly may be driving these differences. It could be related to the spatial patterns of the sample points, as this seems feasible given the very distinct patterns shown in Figure 3, especially given the lack of samples in the east for Landsat_(SceneCloudFree)_. However, we could not dismiss the impacts of other factors such as difference in sample collection methods or size of dataset.

The strength of this study was in the extensive dataset and model validation against in situ measurements, which enhanced the accuracy of the predictions through the entire year across all seasons. However, this study had limitations such as the inherent errors in the Landsat temperature measures due to cloud interference. Cloud cover and smaller lakes also limited coverage in the eastern U.S. for the Landsat_(SceneCloudFree)._ While the Landsat_(LakeCloudFree)_ model showed better performance than the Landsat_(SceneCloudFree)_ model, it consistently underestimated temperature (Fig. 8 and 9). The cloud filtering process applied to Landsat_(SceneCloudFree)_ resulted in a longitudinal split, with warmer error in the western U.S. and cooler error in the eastern U.S. (Fig. 9c). One potential explanation for this pattern was the limited in situ training and validation data, which was constrained by the <1% cloud cover and shadow filter. This filtering reduced the number of usable Landsat_(SceneCloudFree)_ data days, mostly concentrated in summer months. This also limited the longitudinal range, skewed toward the west, and diminished the occurrence of water temperature extremes at both the lower and upper ends of the range. Other potential explanations included water vapor differences related to the Landsat atmospheric correction (Li et al. 2013) or clear sky days skewed toward the west from March through November (Ermida et al. 2019; see thier Figures 3e through 5e). The spatial and temporal gaps in the satellite data due to cloud cover and revisit intervals remains a challenge, requiring models to fill these gaps for higher resolution temporal events such as daily to weekly cyanoHAB dynamics. We did not include wind speed or wind direction, even though wind can affect lake surface temperature through water advection, heat flux, and vertical mixing especially in larger lakes with greater fetch (Bolgrien and Brooks 1992; Laird et al. 2003; Plattner et al. 2006). Shoreline length and surface area may partially account for the impact of wind fetch. The omission of wind may contribute to model error under episodic high-wind conditions and in larger lakes. Future work could evaluate whether adding wind fields improves accuracy.

Additional research will be required to improve the accuracy of the satellite derived surface water temperature measurements. Comparisons between newer machine learning model approaches, such as Long Short-Term Memory and Random Forest, show comparable performance results (Ghosh et al. 2022; He et al. 2019; Li et al. 2022; Rosyad and Maghridlo 2025). Use of hybrid Long Short-Term Memory - Random Forest approaches may improve results, providing an opportunity for future studies. Expansion of the in situ dataset to include more measurements from a range of geographic, seasonal, and anthropogenic conditions will further improve the surface water temperature model performance and representativeness. The Landsat_(LakeCloudFree)_, in situ, and validation datasets capture most of the range of elevation in the NHD Plus waterbodies, with the exception of waterbodies that occur above approximately 3,500 meters, which may be particularly susceptible to temperature changes (Rangwala and Miller 2012). Future studies may also investigate long term impacts of climate change on lake temperature and the subsequent effects on these ecosystems and relation to cyanoHAB occurrence.

For national, management-oriented forecasting, we used a model approach that provided clear assumptions, integrated uncertainty, had lower data requirements, and can be developed and deployed rapidly compared with complex simulation frameworks for ecological management (Cuddington et al. 2013). We note that random forests perform best within the range of observed conditions, and that limited winter or extreme observations may likely result in lower performance. Process-based models are better situated to extrapolate beyond the training data. However, these require rich, long, and spatially expansive datasets to estimate parameters meaningfully. Many of the data sets required for process models are sparse in time and space. Lake sampling methods in the WQP varied by reporting organization and the intensity of sampling varied depending on season and year (Schaeffer et al. 2018b). In addition, information such as stream and river flow would be required, where most of these systems were not gauged. This excluded the option of a purely process based model for our study, but we acknowledge these models provide benefit as they are developed for more systems over the longer term. The Process Guided-Deep Learning models may provide future promise, where Read et al. (2019) tested performance of extrapolation across 68 lakes and found this approach was still limited, similar to our random forest approach. The accuracy was influenced by the number of training observations, performance was best within the range of observed conditions, and the Process Guided-Deep Learning models outperformed when the inputs and training data were higher quality. Temperature is important in regulating biological processes within aquatic ecosystems. The rate of biological growth, including metabolism and photosynthesis, is temperature dependent. Increased temperatures may exacerbate cyanoHAB toxin production (Walls et al. 2018) and be associated with increased stratification and dissolved oxygen depletion (Missaghi et al. 2017; Tan et al. 2018). The 2022 annual temporal trend of model temperature in this study also follows previously published ecological patterns of cyanobacteria seasonal biomass changes (Feng et al. 2018; Marshall and Peters 1989; Rohwer et al. 2023), indicating the strong potential benefit of such temperature models in forecasting cyanoHABs. Temperature also influences physical processes such as lake turnover, stratification, and mixing impacting nutrient cycling and potential for cyanoHABs. Lake temperature changes are also impacted by climate change which may lead to extended growth seasons (Sahoo et al. 2016). Quantifying these temperature dynamics are important for the adaptive management of lakes.

Accurate prediction of lake surface water temperature is important for forecasting cyanoHABs, which can have a range of human, animal, and environmental health effects. Therefore, the implications of this study are relevant for water quality management, ecosystem services, and public health protection. This study highlights some precautions, such as the impact of clouds on remote sensing accuracy, to consider for predictive modeling relevant to lake temperatures. The continuous time series of modeled temperature data can support early warning forecasts for cyanoHABs (Schaeffer et al. 2024).

## Conclusion

This study provides a novel contribution to lake temperature monitoring relevant to cyanoHAB forecasting. Validation of the Landsat derived surface water temperature and development of random forest models demonstrate methods to generate continuous temperature datasets for 2,192 of the largest U.S. lakes that are resolved by Sentinel-3 OLCI. The random forest models using in situ temperature measurements were not biased from cloud presence and demonstrated better model performance when compared models using Landsat surface temperature. This work emphasized the need for accurate and reliable temperature data for cyanoHAB forecasting, where combining in situ data with models fills critical spatio-temporal gaps supporting the ability to not only predict lake surface water temperature but also predict cyanoHAB events in order to protect aquatic ecosystems.

## CRediT authorship contribution statement

Blake A. Schaeffer: Conceptualization, Writing – original draft, Writing – review & editing, Resources, Supervision, Project administration, Funding acquisition. Hannah Ferriby: Conceptualization, Methodology, Software, Formal analysis, Investigation, Writing – original draft. Wilson Salls: Conceptualization, Methodology, Software, Investigation, Writing – original draft. Natalie Reynolds: Methodology, Software, Formal analysis, Investigation, Writing – original draft. Jeff Hollister: Conceptualization, Writing – original draft, Validation, Writing – review & editing. Betty Kreakie: Conceptualization, Writing – original draft, Validation, Writing – review & editing. Stephen Shivers: Writing – original draft, Writing – review & editing. Brent Johnson: Writing – original draft, Writing – review & editing. Olivia Cronin-Golomb: Validation, Writing – review & editing, Kate Meyers: Writing – review & editing, Maxwell Beal: Validation, Writing – review & editing

## Acknowledgements

This material is based upon work supported by the NASA Ocean Biology and Biogeochemistry Program/Applied Sciences Program (proposals 14-SMDUNSOL14-0001 and SMDSS20-0006). Work was also supported by the EPA, and Oak Ridge Institute for Science and Education (ORISE). This article has been reviewed by the Center for Environmental Measurement and Modeling and approved for publication. Mention of trade names or commercial products does not constitute endorsement or recommendation for use by the U.S. Government. The views expressed in this article are those of the authors and do not necessarily reflect the views or policies of the EPA.

